# Temporal transcriptomics and molecular dynamics identify a serine-type endopeptidase as a key regulator of dengue virus infection in *Aedes aegypti*

**DOI:** 10.1101/2025.11.20.689538

**Authors:** Samip Sapkota, Farzana Yeasmin, Soharth Hasnat, Soaibur Rahman, A.D.A Shahinuzzaman, M. Nazmul Hoque

## Abstract

The dengue virus (DENV), a major global pathogen causing over 400 million annual infections, relies on the mosquito *Aedes aegypti* as its primary vector. Intriguingly, *A. aegypti* sustains persistent DENV infection without exhibiting apparent pathology, indicating a highly adapted and regulated host-virus relationship. However, the temporal gene expression dynamics that govern this finely balanced interaction remain poorly understood. We performed a comprehensive transcriptomic analysis using 12 paired-end RNA-seq datasets from gene expression omnibus (GEO; GSE222893), comparing naïve and DENV-infected *A. aegypti* samples at Days 1, 2, and 7 post-infection. A robust bioinformatics pipeline (STAR → FeatureCounts → DESeq2 → g:Profiler) was employed to identify differentially expressed genes (DEGs), explore functional annotations, and resolve temporal patterns via principal component analysis. Finally, a molecular dynamics simulation (MDS) was performed to check the molecular stability of the highly expressed gene. Our temporal analysis identified *LOC5570687*, a gene encoding a serine-type endopeptidase, as the most significantly differentially expressed transcript across all infection time points. Functional annotation confirmed its role in proteolysis, implicating it in the cleavage of flaviviral polyproteins, a critical step in viral replication. Principal component analysis revealed distinct transcriptional divergence at Day 1, immune modulation at Day 2, and convergence by Day 7—marking virion maturation. Downregulation of *LOC5570687* in DENV-exposed mosquitoes was temporally associated with enhanced viral replication, indicating its potential role as a molecular switch between antiviral defense and viral exploitation. A 100 ns MDS was proof of the structural stability, compactness, and dynamic properties of the highly expressed protein. This study uncovers the temporally dynamic transcriptional landscape of *A. aegypti* during DENV infection and the serine-type endopeptidase *LOC5570687* as a critical regulator of viral pathogenesis. These findings provide a molecular framework for understanding vector competence and propose the *LOC5570687* as a promising target for vector-based intervention strategies to disrupt DENV transmission.

## Introduction

Dengue virus (DENV), a member of the Flavivirus family, remains one of the most prevalent mosquito-borne viral pathogens worldwide, infecting an estimated 400 million people annually across over 100 endemic countries [1, 2]. The primary vector of DENV, *A. aegypti*, exhibits remarkable adaptability and competence in facilitating viral transmission [3, 4]. Despite extensive research into the immunological and ecological factors influencing vector capacity, the molecular mechanisms by which *A. aegypti* modulates its gene expression in response to DENV infection remain only partially understood [5]. Mosquitoes, unlike mammalian hosts, do not suffer overt pathogenic effects from DENV infection. Instead, the virus establishes a persistent, non-lethal infection that progresses through key tissues, including the midgut, hemolymph, and salivary glands, ultimately enabling transmission to a new host [6]. This unique host-virus equilibrium raises critical questions about the transcriptional adjustments that support both immune evasion and efficient viral replication [7]. Recent advances in RNA-sequencing (RNA-seq) technologies now enable genome-wide quantification of gene expression, offering a high-resolution view into the temporal dynamics of mosquito antiviral responses [8, 9].

Previous transcriptomic studies have identified innate immune signaling pathways, including the Toll, IMD, and JAK-STAT cascades, as important regulators of antiviral defense in mosquitoes [10, 11]. However, much of the current knowledge is derived from studies lacking temporal resolution or focused on a single infection stage [12]. As DENV replication is a time-sensitive process, a multi-timepoint analysis is essential for disentangling early antiviral responses from viral adaptation mechanisms during infection progression [13]. Today, computational biology is commonly employed to identify drugs, targets, and explore molecular details [14-16]. Current studies encourage the design of research aimed at revealing differentially expressed genes at three post-infection time points.

In this study, we applied a comprehensive bioinformatics pipeline (STAR → FeatureCounts → DESeq2 → g:Profiler) to analyze RNA-seq data from *A. aegypti* mosquitoes infected with DENV at three post-infection time points: Day 1, Day 2, and Day 7. By contrasting DENV-infected and naïve-infected transcriptomes across these intervals, we aimed to () identify differentially expressed genes (DEGs) associated with infection stages, (ii) uncover uncharacterized genes with potential roles in DENV replication or immune modulation, and (iii) map functional enrichment trends underlying host-pathogen interactions. Our findings revealed *LOC5570687*, a previously unannotated gene, as the most significantly differentially expressed across conditions. Functional analysis predicted its role as a serine-type endopeptidase, a class of proteolytic enzymes frequently co-opted by flaviviruses to process viral polyproteins. Moreover, PCA highlighted distinct transcriptional shifts between infected and control groups, indicating that DENV infection induces a stage-specific reprogramming of host gene expression. Together, these results offer novel insights into the temporal biology of DENV-mosquito interactions and identify potential molecular targets for vector-based intervention strategies.

## Methods and materials

### Data collection

Twelve paired-end RNA-seq datasets were retrieved from the Gene Expression Omnibus (GEO) database using the SRA Toolkit [17, 18]. GEO is a public repository that archives and freely distributes high-throughput functional genomic data, including RNA-sequencing experiments. The datasets comprised two experimental conditions: naïve-infected and dengue-infected samples. Specifically, six datasets (SRR23079315, SRR23079316, SRR23079318, SRR23079319, SRR23079321, SRR23079322) corresponded to the naïve-infected group, while the remaining six datasets (SRR23079324, SRR23079325, SRR23079327, SRR23079328, SRR23079330, SRR23079331) represented the dengue-infected. These datasets were selected to enable comparative transcriptomic profiling between naïve and dengue-infected states.

### Conversion, indexing, and mapping of RNA-seq data

The datasets were first converted into FASTQ format using the fasterq-dump command-line utility for each sample [19]. Following conversion, genome indexing was performed using the STAR aligner to prepare a reference genome index stored in a designated directory (dengue.index) [20]. This step utilized the *A. aegypti* reference genome (GCF_002204515.2_AaegL5.0_genomic.fna) and its corresponding annotation file (GCF_002204515.2_AaegL5.0_genomic.gtf), both downloaded from the NCBI RefSeq database [21]. The mapping step involved aligning each of the twelve FASTQ samples individually to the indexed genome using STAR to produce alignment files in the BAM (Binary Alignment Map) format [22]. BAM files are compressed using the BGZF (Blocked GNU Zip Format) and store alignment data in a binary format not readable by humans, unlike SAM (Sequence Alignment Map) files, which contain human-readable, uncompressed alignment information. The resulting BAM files were used for downstream transcriptomic analysis. Reference specifications for the SAM/BAM format were followed as outlined by the official SAMtools [23].

### Gene quantification using feature counts

Gene-level quantification of aligned RNA-seq data was performed using the FeatureCounts tool from the Subread package [24]. FeatureCounts is a highly efficient and widely used program designed to summarize read counts over genomic features, enabling accurate gene expression analysis. In this study, FeatureCounts was applied to the BAM files generated from the STAR mapping step, using the corresponding annotation file (GCF_002204515.2_AaegL5.0_genomic.gtf) to assign mapped reads to specific genomic features. The program was configured to count only uniquely aligned reads and to exclude multi-mapping and low-quality alignments, thereby ensuring the reliability of gene expression estimates. Parameters were adjusted to match the strandedness and layout of the paired-end reads [24]. The output from FeatureCounts provided a matrix of raw read counts for each gene across all 12 RNA-seq samples, which was subsequently used as input for downstream differential gene expression analysis.

### Metadata compilation

A metadata file was constructed using Microsoft Excel to organize and annotate essential information for each of the 12 RNA-seq samples [25]. This metadata included GEO accession IDs, sample names, strain information, organism identity (*A. aegypti*), treatment conditions (naïve or dengue-infected), and associated time points post-infection (Day 1, Day 2, and Day 7). The metadata served as the colData input for downstream DESeq2 analysis, providing the necessary experimental design variables for differential expression modeling. Special attention was given to maintaining row consistency and sample order between the metadata (colData) and the gene count matrix (countData) to ensure compatibility and accurate mapping of sample-level information during statistical analysis [26].

### Differential gene expression analysis

DGE analysis was performed in RStudio using the DESeq2, tidyverse, ggplot2, and apeglm packages [26-28]. The gene count matrix (g.txt) and metadata file (metadata.csv) were imported into RStudio, and a DESeqDataSet object (dds) was created. The analysis included data pre-filtering, normalization, dispersion estimation, and model fitting following standard DESeq2 workflow guidelines[28]. An adjusted p-value threshold of 0.01 was set to identify significantly differentially expressed genes [29]. An epsilon value of 1.0 was used to reduce numerical bias in calculations. Shrinkage of log2 fold changes was applied using the apeglm method, and diagnostic plots were generated using ggplot2 to assess data quality and dispersion [30].

### Data pre-filtering and contrast setup

Prior to differential expression analysis, low-count genes were filtered from the raw count matrix (g.txt) to improve statistical power and reduce noise. Genes with fewer than 10 total reads across all samples were removed, reducing the dataset from 19,269 to 15,307 genes, and the filtered matrix was stored in the DESeq2 object (dds) [31]. For accurate comparison between conditions, releveling of the reference group was performed. By default, R orders factor levels alphabetically, which can affect the interpretation of contrasts. To explicitly define the comparison of interest, naïve-infected samples were set as the reference group, allowing for differential expression analysis of dengue-infected samples relative to naïve controls. This ensured that log2 fold changes were calculated in the biologically meaningful direction (dengue vs. naïve) [31].

### DESeq2 data analysis and log fold change shrinkage

DGS analysis was carried out using the DESeq2 package in Rstudio [31]. The pre-filtered count matrix was passed into the DESeq2 pipeline using the command dds <- DESeq(dds), which sequentially performs normalization, dispersion estimation, and negative binomial generalized linear modeling [32]. During execution, DESeq2 provided progress messages summarizing each step, including estimation of size factors, dispersion values, and the relationship between mean expression and dispersion, followed by model fitting. To enhance interpretability and stabilize variance in genes with low counts, log2 fold change (LFC) shrinkage was applied using the lfcShrink function with the apeglm method [32]. This approach offers more accurate effect size estimates compared to traditional shrinkage methods, improving gene ranking and visualization for downstream analyses.

### Identification of highly significant genes

To identify and visualize highly significant differentially expressed genes (DEGs), bar plots and volcano plots were generated using the results_shrunk_sig dataset in Rstudio [32]. The ggplot2 package was employed to construct bar plots, where genes with the highest log2 fold changes were identified as upregulated based on bar height [27]. The gene names corresponding to the tallest bars were labeled directly above the bars, indicating their strong differential expression. Volcano plots further illustrated the distribution of DEGs based on statistical significance (adjusted p-values) and magnitude of expression changes (log2 fold change) [27, 32]. For comparative analysis, two output files (.tsv format) were generated using the write function: naivevsdengue_RNA_SEQ.tsv (naïve vs. dengue-infected) and dengue1vsnaive_RNA_SEQ.tsv (dengue-infected vs. naïve). Each file contained the top five upregulated and five downregulated genes, which are hypothesized to play important roles in the replication and pathogenesis of DENV in *A. aegypti*.

### Gene ontology analysis and functional annotation

Gene ontology (GO) enrichment analysis was performed using the g:GOSt tool available in the g:Profiler web server (https://biit.cs.ut.ee/gprofiler/) [32]. This tool utilizes Fisher’s one-tailed test to identify statistically significant overlaps between input gene sets and annotated biological terms. A list of significantly DEGs was submitted in the query section, with the significance threshold set to 0.01. The multiple testing correction method was set to Benjamini–Hochberg false discovery rate (FDR) to control for type I error [33]. This analysis facilitated the prediction of biological processes, molecular functions, and cellular components associated with both characterized and uncharacterized genes. To further explore the biological relevance of highly significant genes, functional annotations were retrieved from the NCBI Gene database [34]. Among the top four genes analyzed, two had predicted roles, one had a function confirmed by experimental evidence, and one was classified as a non-coding RNA (ncRNA), remaining functionally uncharacterized due to the lack of protein-coding potential.

### Molecular stability analysis of the protein of a highly expressed gene

Molecular dynamics simulations (MDSs) were performed using the YASARA Structure package to assess the conformational stability, compactness, and dynamic behavior of the target protein [35]. The experimental structure was first prepared by optimizing protonation states at physiological pH (7.4), followed by energy minimization using the YASARA energy minimization protocol, which applies steepest descent and simulated annealing to remove steric clashes and unfavorable contacts. The AMBER14 all-atom force field was employed for both protein and solvent modeling [16]. The simulation system was built by placing the protein in a cubic simulation box with 10–12 Å padding and solvated using TIP3P explicit water molecules. The system was neutralized by adding counter-ions and further adjusted to 0.9% NaCl to mimic physiological ionic strength [36]. Periodic boundary conditions were applied, and long-range electrostatics were calculated using the Particle Mesh Ewald (PME) method. A cutoff radius of 8.0 Å was set for short-range van der Waals and Coulomb interactions. Prior to production, the system underwent a two-step minimization followed by a short equilibration phase while gradually heating the system to 298 K (25°C). The simulation was conducted under an NPT ensemble, maintaining constant temperature using a Berendsen thermostat and constant pressure using a Manometer-based barostat [37]. A time step of 2.0 fs was used, and snapshots were saved at regular intervals for downstream analysis. Production MD was executed for 100 ns without applying any positional restraints.

### Trajectory and structural analysis

Post-simulation trajectory analyses were performed using built-in YASARA analysis macros. Structural stability was assessed by calculating root mean square deviation (RMSD) of Cα atoms, while protein compactness was measured using radius of gyration (Rg). Total potential energy, non-bonded interaction energy, and hydrogen bond counts were monitored to evaluate energetic stability and proper system equilibration. Secondary structure content was quantified using the YASARA DSSP-based module across the full simulation time. To investigate residue-level flexibility, root mean square fluctuations (RMSF) were calculated for backbone and side-chain atoms [33]. Residue-residue contact maps, contact numbers, and cross-correlation analyses were carried out to evaluate tertiary packing consistency and dynamic network preservation across the trajectory.

## Results

### Analysis of raw and normalized counts in dengue and naïve infection samples

The comparison between raw and normalized log2-transformed RNA-seq counts revealed a significant difference in data distribution uniformity across samples (**Fig. 1**). The x-axis represents the log2-transformed counts, while the y-axis displays individual sample identifiers (SRR23079315 through SRR23079331). Blue boxes correspond to raw counts, and red boxes correspond to normalized counts. Each box plot shows the distribution of counts per sample, with the black vertical lines indicating the median (central tendency), and the whiskers representing variability beyond the interquartile range (IQR) (**Fig. 1**). The raw counts exhibit minimal differences in median location and IQR width across samples. In contrast, the normalized counts show slightly better alignment of medians and a more uniform IQR width, as expected, since normalization aims to make the count distributions as similar as possible. However, the initial variability in the raw counts was already low, indicating the samples had similar overall sequencing depth. The normalization confirmed and slightly enhanced this uniformity. This adjustment ensures reliable downstream analysis, facilitating the accurate detection of differentially expressed genes by improving comparability across samples (**Fig. 1**).

**Fig 1.**
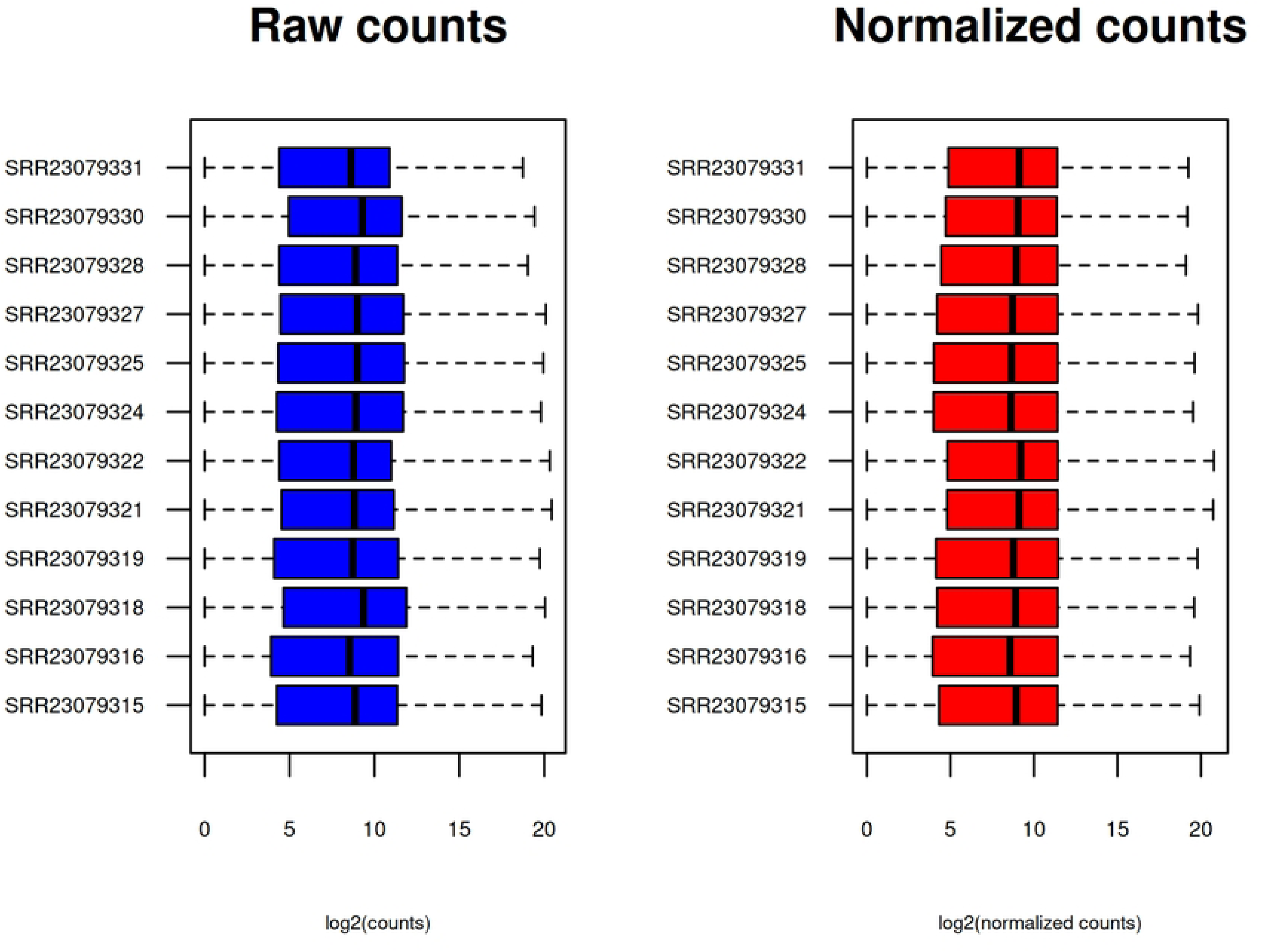
The divergence between raw counts and the normalized counts in dengue and naïve infection samples. The graph amplifies the fluctuations in the raw counts while reducing the volatility in the normalized counts.

### Comparison of dispersion estimates between naïve and dengue-infected samples

To assess the similarity and reliability of gene expression variability across naïve and DENV-infected samples, dispersion estimates were computed using the DESeq2 statistical framework, which models variance as a function of mean normalized counts. Dispersion represents the variability of count data and is inversely related to expression abundance, meaning that genes with low mean normalized counts generally exhibit higher dispersion and greater expression volatility, whereas highly expressed genes tend to show lower dispersion and more stable expression profiles. The initial dispersion estimates displayed the expected spread for low-count genes, while the fitted trend line demonstrated a gradual decline in dispersion with increasing mean values, reflecting a biologically consistent distribution (**Fig. 2**). Moreover, the final dispersion estimates closely followed the fitted curve, indicating that the model successfully captured the underlying variance structure for both naïve and dengue-infected datasets. The agreement between observed dispersion values and the fitted trend confirms accurate variance estimation, robust normalization performance, and appropriate statistical modeling. Collectively, these findings validate that gene expression variability was reliably characterized, thereby providing a sound statistical basis for subsequent differential expression analyses (**Fig. 2**).

**Fig. 2.**
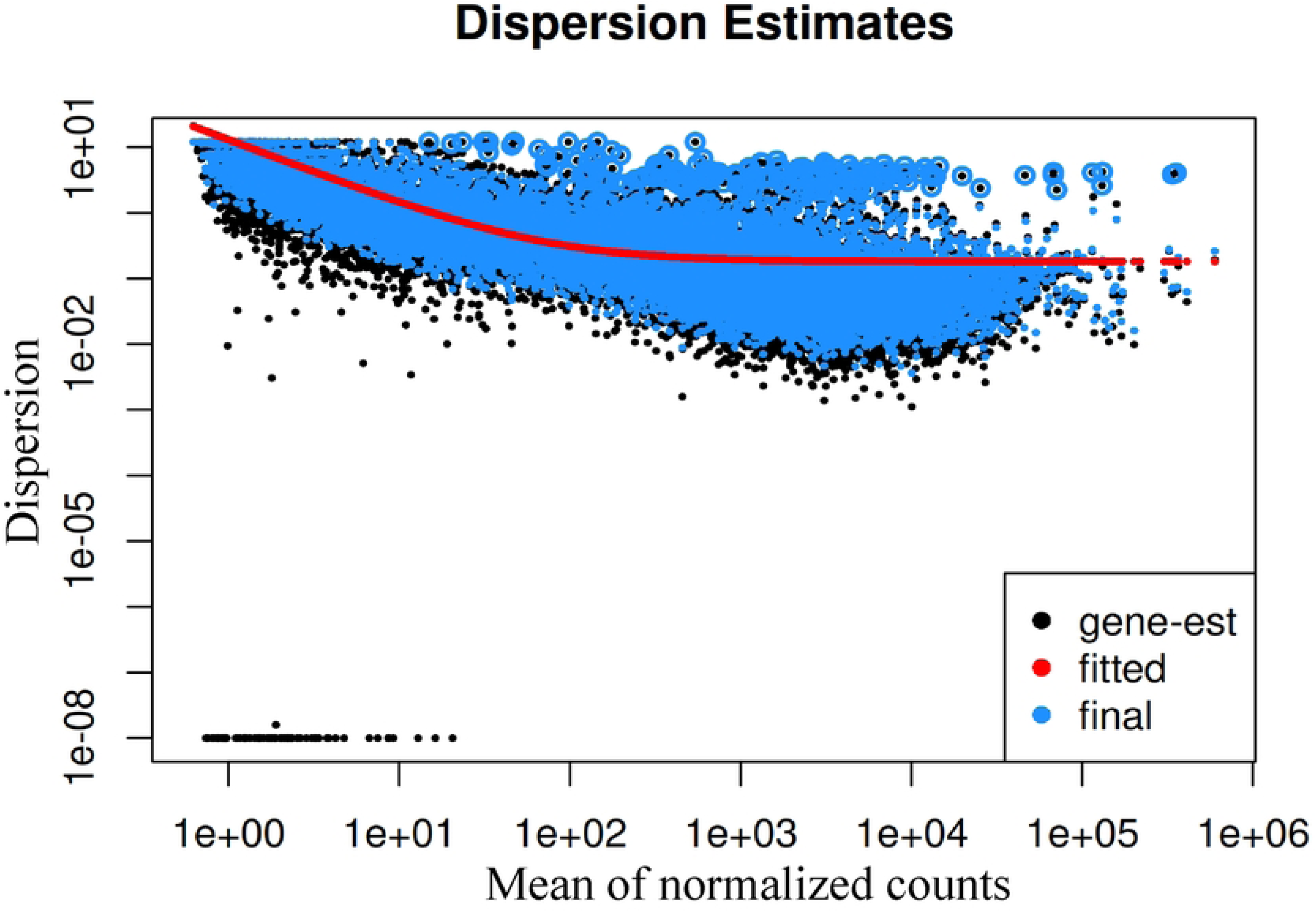
Dispersion estimate plot correlating the naïve infection and dengue infection. The X-axis illustrates the mean of normalized counts, while the Y-axis denotes dispersion. The black dots indicate the initial dispersion estimate. The red trend line represents the dispersion recession procedure with an increasing mean of normalized counts. The blue circles reflect the final dispersion rate.

### Interpretation of mean–average plots comparing gene expression between naïve and dengue infection

To characterize differential gene expression between naïve and dengue-infected mosquito samples, mean–average (MA) plots were generated (**Figs. 3** and **4**) to illustrate the relationship between log fold change and mean normalized counts. Each point in the plots represents an individual gene, where grey points indicate genes with minimal expression differences, while blue points denote significantly differentially expressed genes. The horizontal line at log fold change = 0 represents no change in expression, and the red boundary lines define significance thresholds. In **Fig. 3** (dengue-infected vs. naïve), blue points below the red threshold indicate genes downregulated in dengue-infected samples, whereas in **Fig. 4** (naïve vs. dengue-infected), blue points above the threshold indicate genes upregulated in naïve samples. Together, these results demonstrate clear transcriptional divergence between conditions and identify distinct sets of significantly regulated genes, offering valuable insights into infection-associated molecular responses and potential biomarker candidates for future research.

**Fig. 3.**
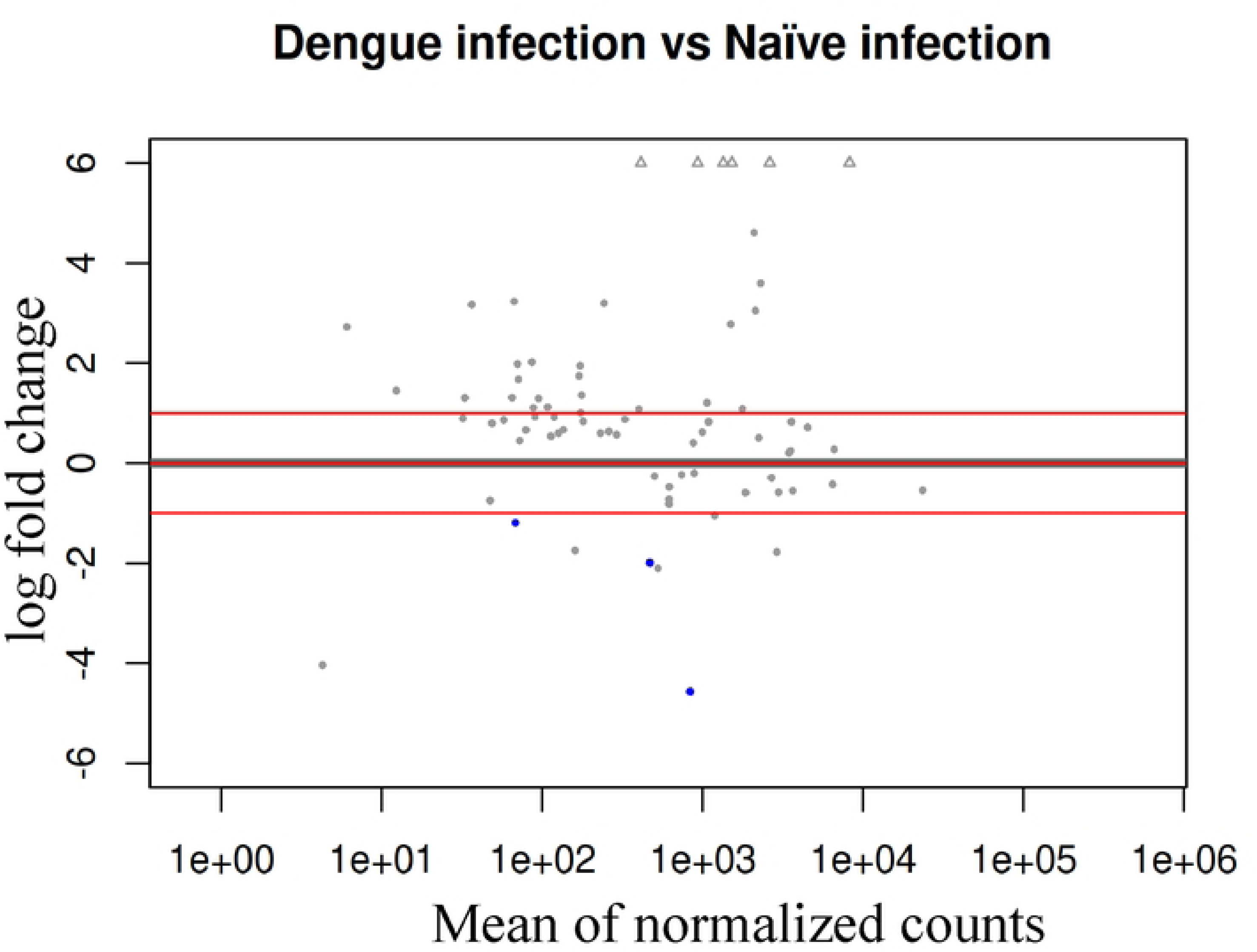
The mean–average (MA) plot represents log fold change and the mean of normalized counts on the X and Y axes, respectively. The graph illustrates the downregulated genes shown as blue dots below the red line in the negative log fold change area. These downregulated genes are present in mosquitoes infected with DENV compared to naive infection mosquitoes.

**Fig. 4.**
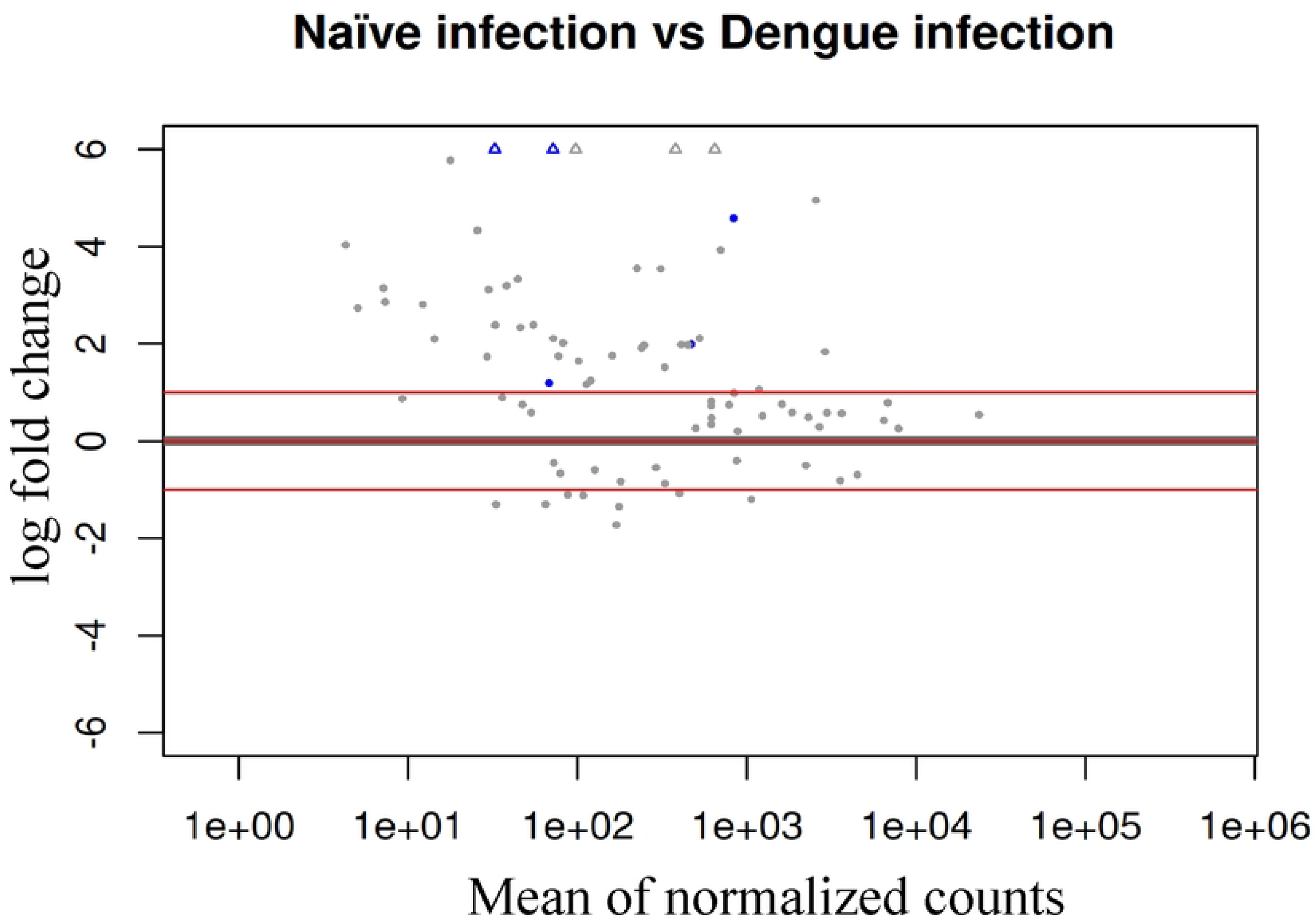
Mean–average (MA) plot representing log fold change and mean of normalized counts on the X and Y axes, respectively. The graph illustrates the up-regulated genes shown as blue dots above the red line, indicating positive log fold change. The up-regulated genes are present in naïve infection versus DENV infection mosquitoes.

### Expression profile of the most highly upregulated gene across naïve and dengue-infected samples

Transcriptomic analysis of *A. aegypti* samples infected with DENV versus naïve controls revealed *LOC5570687* as the most significantly upregulated DEG. As illustrated in **Fig. 5**, this uncharacterized gene (NCBI accession: *LOC5570687*) exhibited pronounced expression variability across biological replicates. Notably, its peak expression occurred in sample SRR23079319 (Naïve-infected, Day 2, Replicate 2; GSE222893), suggesting potential baseline regulatory functions in uninfected mosquitoes. This aligns with prior evidence implicating *LOC5570687* in DENV replication mechanisms [38]. The gene’s consistent upregulation in multiple DENV-exposed samples (e.g., SRR23079315, SRR23079324) further underscores its significance in mosquito antiviral responses (**Fig. 5**).

**Fig. 5.**
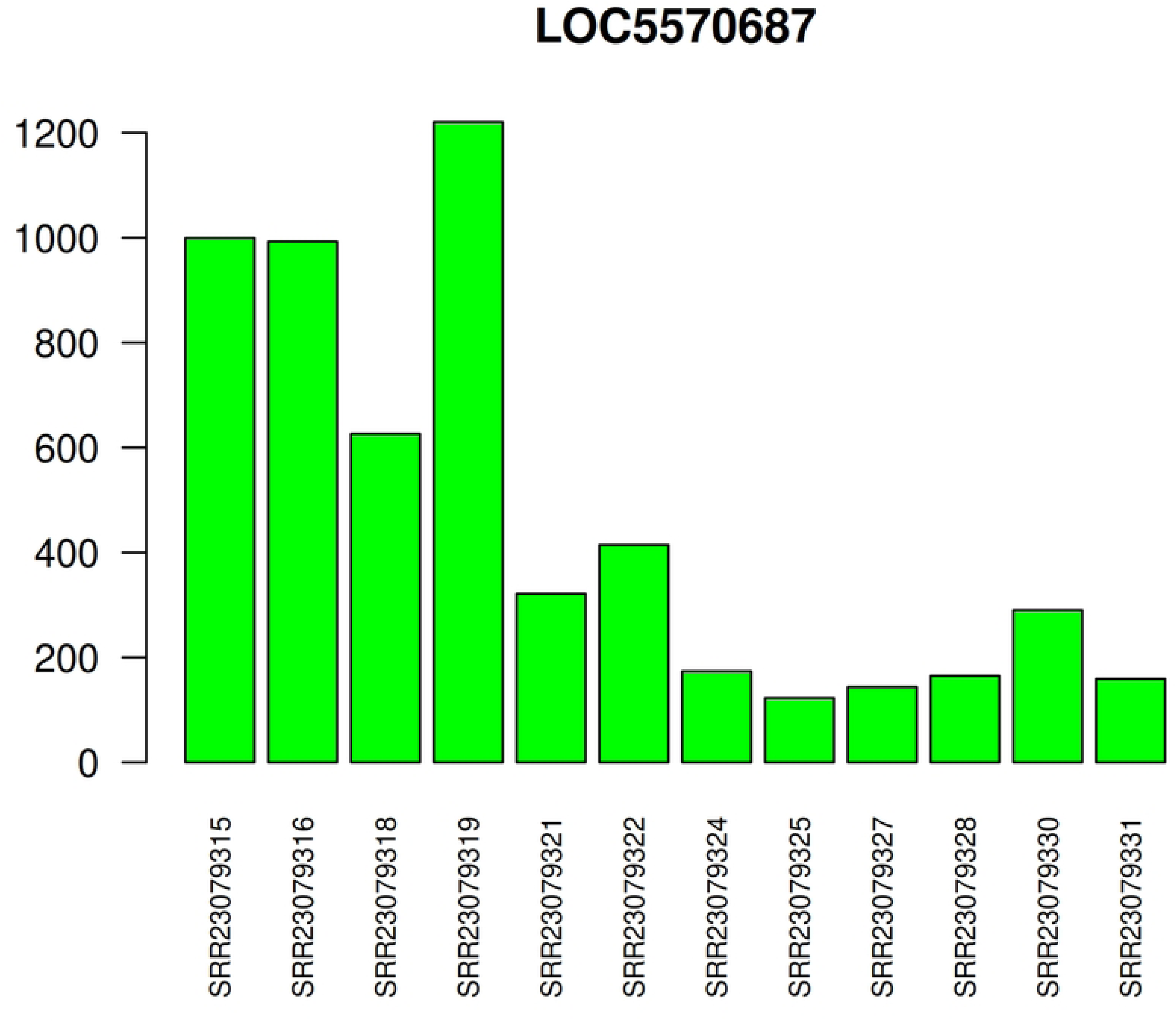
Expression profile of *LOC5570687* in *A. aegypti* during Dengue infection vs. naïve controls. Bar plot displays normalized expression levels of the top differentially expressed gene (*LOC5570687*) across 12 samples. Samples include Dengue-infected (DENV) and naïve-infected mosquitoes at Day 2 post-infection (biological replicates 1–3). Peak expression occurred in the naïve sample SRR23079319. Gene accession: *LOC5570687*; Data source: GSE222893.

### Functional characterization of *LOC5570687* reveals serine-type endopeptidase activity in DENV replication

Gene Ontology (GO) analysis was performed using g:Profiler to functionally characterize the uncharacterized gene *LOC5570687*, identified as the most significantly differentially expressed gene in DENV-infected *A. aegypti*. Under stringent parameters (significance threshold: *p* < 0.01, Benjamini-Hochberg correction), enrichment analysis identified serine-type endopeptidase activity (GO:0004252) as its primary molecular function. This protease function predicts a critical role in viral pathogenesis: upon ingestion of DENV by naïve mosquitoes, *LOC5570687* may cleave viral polyproteins at specific serine residues to generate structural and non-structural viral components essential for replication (**File S1**). This proteolytic activation mechanism aligns with established models of flavivirus processing and underscores the gene’s potential as a regulatory switch in early infection stages.

### PCA highlights temporal divergence in *A. aegypti* responses to DENV infection

PCA of *A. aegypti* transcriptomes revealed distinct temporal shifts in host responses to dengue infection (**Fig. 6**). The PCA plot (70% variance explained; PC1: 52%, PC2: 18%) showed clear separation between dengue-exposed and naïve-infected mosquitoes across Days 1, 2, and 7 post-infection. At Day 1, dengue-exposed mosquitoes (pink) diverged significantly from naïve counterparts (yellow), indicating early viral engagement. In naïve mosquitoes, dengue entered midgut cells via receptor-mediated endocytosis, activating *LOC5570687*-linked serine-type endopeptidase activity to process viral polyproteins. By Day 2, naïve mosquitoes (cyan) maintained high *LOC5570687* expression despite slowed viral replication due to immune defenses. Dengue-exposed mosquitoes (green) showed reduced expression, favoring viral escape and adaptation. By Day 7, both groups converged in PCA space (blue and purple), reflecting similar gene expression as virions matured and invaded salivary glands (**Fig. 6)**. Persistent *LOC5570687* downregulation in dengue-exposed mosquitoes enhanced polyprotein processing, accelerating infection severity.

**Fig. 6.**
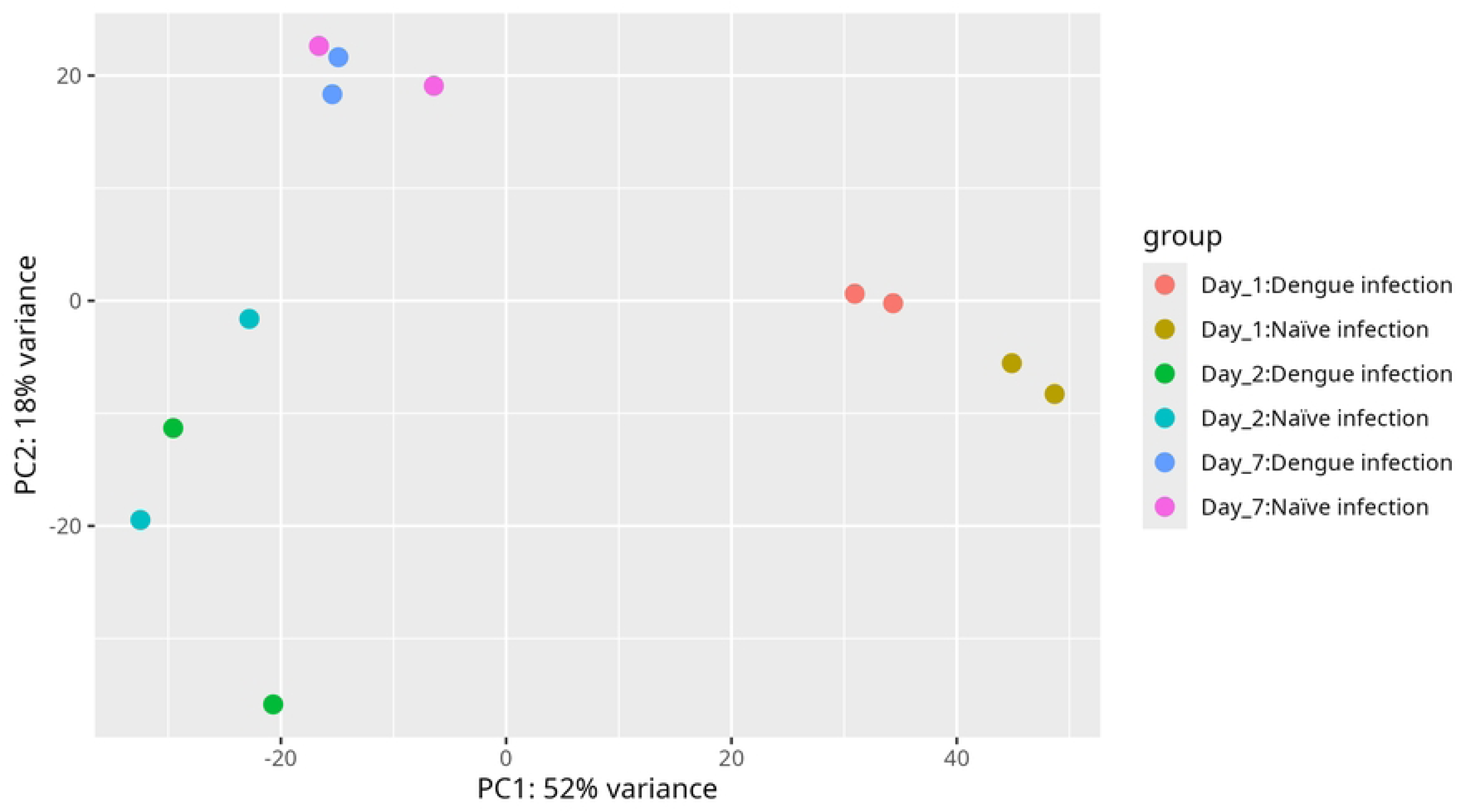
Principal component analysis (PCA) of *A. aegypti* transcriptomes during dengue infection. Scatter plot (PC1: 52%, PC2: 18%; total variance: 70%) shows temporal dynamics in mosquito responses to dengue virus. Distinct clustering between dengue-exposed (pink, green, blue) and naïve-infected (yellow, cyan, purple) groups at Day 1 indicates early transcriptional divergence. By Day 7, convergence reflects similar gene expression patterns associated with virion maturation. Key differentiator: stage-specific regulation of serine endopeptidase gene *LOC5570687* (see text for details). Data source: GEO accession GSE222893.

### Molecular dynamics simulation reveals the stability of highly expressed proteins

Comprehensive MDS were conducted for 100 ns to investigate the structural stability, compactness, and dynamic properties of the protein. The analysis of secondary structure composition (**Fig. 7, Figs. S1–S2**) revealed that the protein preserved its native β-rich fold throughout the simulation. The fractional population of structural elements remained consistent, with β-strands and bends dominating (∼0.30–0.40), accompanied by stable α-helical content (∼0.15–0.18). Minor fluctuations were observed in turns and coil regions (<0.05), primarily localized to surface loops and termini, reflecting natural flexibility rather than conformational disruption. The rate of secondary structure change remained low and stable, confirming structural equilibrium and dynamic steadiness during the simulation. Structural stability was further supported by global parameters (**Figs. S3A–F**). The RMSD of Cα atoms increased during the initial relaxation phase, reaching ∼7 Å and stabilizing after 60 ns, suggesting convergence toward equilibrium. The Rg fluctuated around 26–28 Å, indicating consistent compactness with a brief transient expansion around 60 ns, followed by re-compaction. The total potential energy, non-bonded interaction energy, and hydrogen bond profiles remained stable throughout the production phase, validating proper thermodynamic control and absence of energetic drift. The constant box dimensions confirmed solvent density stability and overall system equilibration. Contact-based analyses (**Figs. S4A–D**) revealed consistent tertiary packing and residue-residue interactions. The total contact number stabilized after equilibration, and the average contact distribution showed that most residues maintained 4–8 atomic contacts, characteristic of a well-folded protein. Low per-residue contact fluctuations and a strong correlation (r = 0.824) between early and late trajectory halves indicated that the intramolecular contact network was conserved, signifying a robust and stable fold. Residue-level flexibility analysis (**Fig. S5**) provided finer insights into dynamic motion. RMSF profiles showed that most residues fluctuated within 2–4 Å, suggesting restricted motion and overall rigidity. Higher RMSF peaks were confined to loop and terminal regions, consistent with inherent local flexibility. The similarity between backbone and side-chain RMSF profiles demonstrated coherent motion without structural distortion, while the smoothed RMSF trends emphasized the dominance of stable, low-flexibility regions across the structure (**Fig. S5**).

**Fig. 7.**
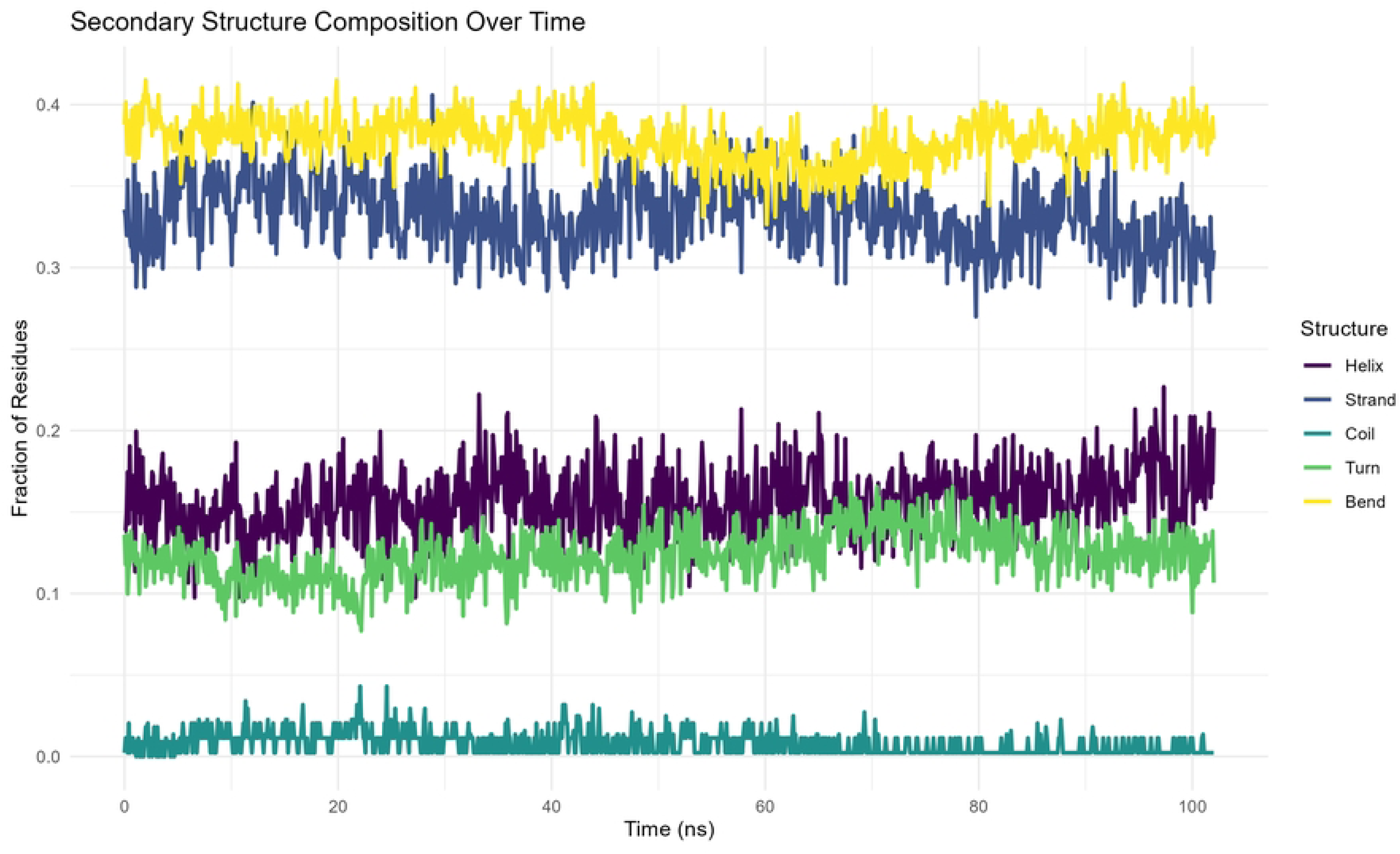
Secondary structure composition over time. The analysis of secondary structure composition throughout the 100-ns molecular dynamics simulation reveals that the protein maintains a stable overall fold with minimal conformational drift. The fractional populations of structural elements remain consistent, with bends and β-strands dominating (≈0.38–0.40 and 0.30–0.35, respectively), reflecting a predominantly β-rich architecture complemented by flexible loop regions. The α-helical content remains steady (≈0.15–0.18), while turns fluctuate slightly (≈0.10–0.13) and coils remain scarce (<0.05), indicating limited disorder. Together, these trends confirm that the protein’s secondary structure is well preserved, suggesting robust structural stability under the simulated conditions.

## Discussion

This study employed an integrated transcriptomic and computational strategy to elucidate the molecular responses of Aedes aegypti to DENV infection across three critical post-infection stages (Days 1, 2, and 7). Through robust differential expression analysis, we identified *LOC5570687* as the most significantly dysregulated gene, revealing its potential role as a serine-type endopeptidase, a protease class commonly exploited by flaviviruses for polyprotein processing. Such host-derived enzymatic assistance is consistent with previous observations where viruses co-opt host serine proteases to optimize replication and maturation of viral structural and non-structural proteins. These findings suggest that the serine-type endopeptidase *LOC5570687* may function as a regulatory molecular switch, modulating antiviral defense under naïve conditions and being repurposed to enhance viral propagation once infection is established. Temporal evaluation of transcriptome changes provided key mechanistic insight into dynamic host-virus interplay [39, 40]. PCA demonstrated clear transcriptomic divergence at Day 1, reflecting early cellular engagement with DENV. At Day 2, naïve samples retained elevated *LOC5570687* expression, implying active host defensive programming, whereas infection-associated suppression of this gene likely facilitated viral immune escape and midgut barrier traversal [41]. By Day 7, expression profiles between groups converged, representing late-stage systemic dissemination and virion maturation [42]. These observations reinforce the concept of time-dependent molecular remodeling, where DENV rewires host protease expression to create a pro-replicative cellular environment [43]. To complement the transcriptomic findings and assess whether the identified protein is structurally competent and potentially functional under physiological conditions, we performed an extensive 100-ns MD simulation [33]. The protein retained its β-rich architecture and secondary structure fractions with minimal deviations, indicating true structural resilience rather than transient conformational persistence. The stability of RMSD, Rg, hydrogen bonding, and energy profiles confirmed that the protein achieved thermodynamic and conformational equilibrium without signs of unfolding or aggregation [44]. Likewise, contact-based metrics demonstrated sustained tertiary packing, while moderate RMSF fluctuations localized to loops and terminal residues aligned with expected functional plasticity. Together, these computational biophysical results suggest that *LOC5570687* encodes a structurally stable, catalytically plausible protein, reinforcing its potential biological relevance during infection and supporting its classification as a viable molecular target. The integration of transcriptomic and molecular dynamics evidence underscores the possibility that DENV modulates host protease biology to create optimal replication conditions, particularly during early and mid-infection phases [45]. From a vector-control standpoint, this study highlights *LOC5570687* as a promising intervention candidate. Gene silencing strategies (e.g., RNAi, CRISPR-Cas9, antisense oligonucleotides) or small-molecule protease inhibitors might impair viral propagation within the vector without compromising mosquito viability—an ideal strategy for reducing transmission without ecological disruption [46, 47]. Despite these advances, experimental validation is necessary to confirm protease activity, substrate specificity, and interaction with DENV polyproteins. Functional knockdown assays, proteolytic cleavage assays, and structural docking with DENV polyprotein intermediates would provide crucial mechanistic clarity [48]. Additionally, incorporating intermediate time-points and field-derived mosquito samples may extend ecological and evolutionary interpretability [49]. In summary, our multi-layered approach reveals that Aedes aegypti displays a temporally dynamic transcriptional response to DENV infection and identifies *LOC5570687* as a structurally stable protease-like target with dual antiviral and pro-viral potential. These findings not only advance understanding of mosquito vector competence but also pave the way for next-generation molecular intervention strategies. Future efforts should focus on the functional characterization of key DEGs and validating intervention approaches in ecologically relevant mosquito populations.

## Conclusion

This study reveals that *A. aegypti* mounts a temporally dynamic transcriptomic response to DENV infection, with distinct gene expression patterns observed across Days 1, 2, and 7 post-infection. The uncharacterized gene *LOC5570687* emerged as a key regulator, exhibiting serine-type endopeptidase activity likely involved in DENV polyprotein processing. Its expression profile suggests a dual role supporting antiviral defense in naïve mosquitoes and facilitating viral replication in infected hosts. These findings underscore the ability of the DENV to manipulate host protease expression to optimize infection and transmission. The rigorous bioinformatic pipeline applied here ensures the robustness of results and offers a framework for studying other vector-borne diseases. Importantly, *LOC5570687* represents a promising target for novel vector-based control strategies. Future research should prioritize experimental validation and assess the relevance of these findings across diverse mosquito populations and DENV serotypes to enhance translational impact.

## Supporting Information

**Fig. S1.** Residue-wise secondary structure evolution over time. The temporal secondary structure map shows that the protein maintains a stable fold throughout the 100-ns molecular dynamics simulation. α-helical and β-strand regions remain highly persistent, indicating structural rigidity within the core, whereas minor transitions among turns, bends, and coils are primarily observed in loop and terminal regions, reflecting localized flexibility rather than unfolding. Overall, the consistent secondary structure pattern supports a dynamically equilibrated and structurally stable conformation under the simulated conditions.

**Fig. S2.** Rate of secondary structure change over time. The temporal evolution of secondary structure dynamics was assessed by monitoring the fraction of residues undergoing structural transitions during the 100-ns molecular dynamics simulation. Moderate fluctuations were observed, with the fraction of changing residues generally ranging between 0.05 and 0.20. Despite these variations, the smoothed trend line remained relatively stable across the trajectory, indicating the absence of major unfolding events or large-scale structural rearrangements. The observed changes likely reflect transient, localized adjustments within flexible loop or coil regions. A slight dip around 50–70 ns may correspond to a brief stabilization period, after which the system returned to a dynamic steady state. Overall, these results confirm that the protein retained its secondary structure integrity under the simulated conditions.

**Fig. S3.** Structural and energetic stability analysis over the 100-ns molecular dynamics simulation. (A) Root-mean-square deviation (RMSD) of Cα atoms: RMSD initially increases from ∼1 Å to ∼6–7 Å within the first 10 ns, indicating structural relaxation. Moderate fluctuations (∼3–5 Å) are observed between 10–55 ns, followed by a rise to ∼7–8 Å around 60 ns, suggesting a conformational adjustment. RMSD then stabilizes near 7 Å, indicating attainment of a new equilibrium state without unfolding. (B) Radius of gyration (Rg): Rg fluctuates around ∼27–28 Å during the equilibration phase, briefly increases to ∼30 Å between 55–65 ns, and then stabilizes near ∼26 Å, suggesting enhanced compactness and attainment of a stable, energetically favorable conformation. (C) Total energy: Total potential energy remains steady throughout the simulation, fluctuating minimally around −2.0 × 10⁶ kJ/mol after equilibration, confirming thermodynamic stability and proper system parameterization. (D) Non-bonded interaction energies: van der Waals and Coulombic interaction energies remain smooth and stable, with electrostatic interactions being the dominant contributor, supporting persistent non-covalent interactions and indicating overall simulation convergence and structural stability. (E) Hydrogen bonds: The number of internal (protein–protein) hydrogen bonds remain stable, fluctuating within the range of ∼250–300 throughout the simulation, indicating preservation of secondary and tertiary structure. Protein–solvent hydrogen bonds show narrow fluctuations around ∼850–900, reflecting stable hydration shell dynamics. These small variations represent normal thermal motion rather than structural destabilization. (F) Periodic box dimensions: The X, Y, and Z dimensions of the simulation box remain constant over time, confirming system equilibration, stable solvent density, and a consistent simulation environment suitable for reliable analysis of protein dynamics.

**Fig. S4.** Protein stability assessment based on residue–residue contact analyses during the 100-ns molecular dynamics simulation. (A) Total atomic contact stability over time: The instantaneous number of residue–residue atomic contacts (red and blue fluctuating lines) and their cumulative averages (red and blue straight lines) rapidly converge to stable plateaus with minimal variation, demonstrating preservation of tertiary packing and hydrophobic core integrity. The stable profiles confirm that the protein maintains a consistent number of internal non-covalent interactions, indicating a well-equilibrated, structurally robust fold without evidence of unfolding or collapse. (B) Average residue–residue contact distribution: Mean atomic contacts per residue (red bars), with the dashed blue line showing the global average, reveal that most residues maintain ∼4–8 contacts, consistent with stable intramolecular packing. Residues near the N-terminus show higher contact frequencies, suggesting localized dense packing or possible interaction interfaces. (C) Contact fluctuation per residue: Standard deviation values (purple bars), with a dashed orange average line, show low fluctuation (∼2–3) for most residues, indicating stable interaction networks and limited local flexibility. Increased fluctuation at the N-terminus reflects expected dynamic mobility due to reduced structural constraints. (D) Contact conservation analysis: Comparison of mean contacts between the first and second halves of the trajectory shows a strong positive correlation (r = 0.824), confirming highly conserved residue-level interaction patterns and persistent tertiary structure stability with only minor local rearrangements.

**Fig. S5.** RMSF-Based Flexibility and Stability Assessment of the Protein during MD Simulation. (A) Backbone Flexibility (Cα Atoms), the RMSF profile of Cα atoms illustrates residue-wise backbone flexibility throughout the simulation. Most residues display RMSF values between 2–4 Å, indicating overall structural stability with limited fluctuation. A few peaks correspond to loop and terminal regions, which are inherently more flexible due to reduced secondary structure constraints. The general trend suggests that the protein maintains a stable backbone conformation during the molecular dynamics trajectory, reflecting a well-preserved structural core. (B) Backbone vs. Side Chain Flexibility: This comparison between backbone and side chain RMSF values reveals that side chains exhibit slightly higher fluctuations than backbone atoms, consistent with their greater conformational freedom. Both profiles follow similar patterns across the residue index, suggesting that local backbone movements are closely associated with side chain dynamics. The absence of major divergence between the two curves implies that no large-scale unfolding or domain movement occurred, reinforcing the overall conformational stability of the protein. (C) Distribution of Backbone Flexibility: The histogram of RMSF values represents the distribution of backbone flexibility across all residues. The data show a predominant clustering around the mean RMSF of 3.43 Å, indicating moderate fluctuations for most residues. This narrow distribution signifies those large deviations are rare, and flexibility is relatively uniform throughout the protein. The shape of the distribution supports the inference that the structure remains compact and stable during the simulation, with minimal high-mobility outliers. (D) Flexibility Trends (5-Residue Moving Average): The smoothed RMSF profile using a 5-residue moving average highlights broader flexibility trends by reducing short-term noise. This representation makes it easier to identify flexible regions and stable structural segments. The overall pattern shows periodic moderate fluctuations interspersed with small peaks, likely corresponding to loop regions or solvent-exposed residues. The absence of persistent high RMSF regions indicates that the protein retained its structural integrity throughout the simulation period.

**File S1.** Viral polyproteins at specific serine residues to generate structural and non-structural viral components essential for replication.

## Data availability

The article contains the data utilized to support the results of the *in-silico* study.

## Author contribution statement

SS, FY, SH, and MNH: conceptualized and designed the study; SS, FY, SH, and AS performed the experiment, acquired and analyzed data, interpreted results, and drafted the original manuscript; AS and MNH critically reviewed and edited the manuscript. Finally, all authors read, edited and approved the manuscript.

## Funding

This *in-silico* research was conducted without financial support from any donor agency or organization.

## Ethical statement

This *in-silico* computational study did not involve human subjects and therefore did not require ethical approval. Furthermore, the authors declare that this manuscript, submitted to PLOS Neglected Tropical Diseases, has been prepared with full adherence to responsible research practices and in accordance with the guidelines of publication ethics.

## Competing interests

The authors have declared that no competing interests exist.

## Consent for Publication

Not applicable

## Notes

### Competing Interest Statement

The authors have declared no competing interest.

